# The pathogenic coronatine toxin hijacks host redox signalling to suppress plant immunity

**DOI:** 10.1101/2023.12.01.569535

**Authors:** Lucas Frungillo, Nodoka Oka, Sang-Uk Lee, Mika Nomoto, Byung-Wook Yun, Yasuomi Tada, Steven H. Spoel

## Abstract

Reciprocal antagonism between the hormones salicylic acid (SA) and jasmonic acid (JA) is particularly important during infection by the bacterial pathogen *Pseudomonas syringae*. *P. syringae* secretion of the virulence-promoting toxin, coronatine, is thought to suppress SA-induced immune responses by manipulating host JA signalling. Here, we report an unexpected JA-independent role of coronatine in promoting pathogen virulence. While JA induced resistance to *P. syringae*, coronatine promoted pathogen virulence by suppressing cellular accumulation of glutathione, a vital antioxidant required for immunity. Coronatine-mediated suppression of glutathione levels prevented activation of NPR1, a redox-sensitive master regulator of SA-responsive immune genes. Moreover, the accrual of nitric oxide (NO) restored virulence of coronatine-deficient *P. syringae*, but was counteracted by expression of the host *S*-nitrosothiol reductase, Thioredoxin *h*5. Thus, our findings indicate *P. syringae* utilises coronatine to suppresses host immunity by precisely manipulating glutathione- and NO-mediated redox signalling networks.

## INTRODUCTION

During incompatible plant-microbe interactions, a burst in production of reactive oxygen and nitrogen intermediates (ROS/RNS) in hosts coordinates specificity and amplitude of defence responses (Yun et al., 2011, 2016). Particularly, the redox active molecule, nitric oxide (NO), is extensively involved in shaping immunity (Kneeshaw et al., 2014; Salgado et al., 2013; Spoel and Dong, 2012). NO regulates protein dynamics mainly through the thiol post-translational modification, *S-* nitrosylation (also known as *S-*nitrosation). *S-*nitrosylation is the covalent attachment of a NO moiety to a reactive thiol group in proteins, to form a protein *S-*nitrosothiol (protein-SNO) (Kovacs and Lindermayr, 2013; Radi, 2013; Yu et al., 2014). Similarly, NO may react with the major antioxidant in cells, glutathione (GSH), to form S-nitrosoglutathione (GSNO), an adduct also capable of generating protein-SNO. Interestingly, evidence indicate that protein-SNO derived from GSNO and NO have distinctive roles in immunity (Yun et al., 2016), raising questions whether hosts and attackers evolved strategies to tune protein-SNO signalling during incompatible interactions.

In plants, protein-SNO levels are controlled through the activity of the protein-SNO reductases, *S-*nitrosoglutathione reductase (GSNOR1) and Thioredoxin-*h*5 (TRX*h*5) (Feechan et al., 2005; Frungillo et al., 2014; Kneeshaw et al., 2014). While GSNOR1 reduces the cellular pool of GSNO, allowing GSNOR1 to indirectly control cellular levels of protein-SNO, TRX*h*5 was shown to directly and selectively reduce protein-SNO (Kneeshaw et al., 2014). Similarly to NO-overproducing *nox1* plants, *gsnor1* mutants exhibit constitutively elevated levels of protein-SNO, which are associated with compromised immune responses to the biotrophic pathogen *Pseudomonas* (Feechan et al., 2005; Kneeshaw et al., 2014; Yun et al., 2011, 2016). Interestingly, overexpression of TRX*h*5 in *nox1* mutants, but not in *gsnor1*, restores protein-SNO levels and immunity to pathogenic *Pseudomonas syringae* (Kneeshaw et al., 2014). These data show that GSNOR1 and TRX*h*5 discriminate between specific groups of protein-SNO (Kneeshaw et al., 2014).

While GSNOR1 and TRX*h*5 provide timely removal of protein-SNO, little is known about the mechanisms underlying specificity in formation of protein-SNO. Here, we provide genetic and biochemical evidence that redox intermediates are at the centre of a molecular battle between *Pseudomonas* and Arabidopsis to control redox signalling in hosts. Our findings imply that pathogens discriminate between different RNS intermediates during host immunity and target specificity in redox-mediated protein post-translational modifications to promote disease.

## RESULTS

### The structural analogues JA and Coronatine induce contrasting immune outcomes

In plants, interplay between the hormones salicylic acid (SA) and jasmonic acid (JA) fine-tunes immune responses (Gimenez-Ibanez et al., 2016; Nomoto et al., 2021; Spoel et al., 2007). Plant immune responses against the hemibiotrophic bacterial pathogen *Pseudomonas syringae* (Psm) is largely mediated by SA (Ellis et al., 2002; Tsuda et al., 2009). Interestingly, despite evidence that JA provides robustness to the plant immune hormonal network (Tsuda et al., 2009), several strains of *Pseudomonas* secrete the virulence-promoting toxin, coronatine, which is thought to suppress SA-induced immune responses by structurally mimicking JA (Feys et al., 1994; Gimenez-Ibanez et al., 2016). Thus, we considered that JA and COR have different roles in plant immunity.

First, to investigate if COR requires the host JA transcriptional machinery to promote disease, we assessed immunity in JA-compromised *myc2,3,4* mutants. The MYC family of transcriptional activators are critical components of JA transcriptional machinery in plants and, consequently, *myc2,3,4* mutant plants fail to reprogram their transcriptome in response to JA (Fernández-Calvo et al., 2011; Zhu et al., 2015). As expected, when infected with the pathogenic, coronatine-producing *Pseudomonas maculicola* ES4326 (*Psm*), *myc2,3,4* mutants displayed enhanced disease resistance compared to wild-type plants (Fig. 1A). Next, we examined the impact of JA and COR on transcriptional changes during immunity. To that end, we assessed transcriptional changes in wild-type plants after leaf immunization with SA in the presence or not of JA or COR. Leaf spray with SA induced strong transcriptional activation of SA-responsive gene *PR1* when compared to mock-treated plants, which was largely insensitive to JA or COR co-treatment (Fig. 1B). While SA treatment in the presence of JA or COR markedly activated transcription of JA-synthetic gene *LOX2* compared to SA and mock treatments, JA and COR selectively induced transcriptional activation of JA-responsive genes *PDF1.2* and *VSP2* (Fig. 1B), respectively. Taken together, these data indicate that JA and COR differentially regulate the plant immune hormonal network.

**Figure 1.**
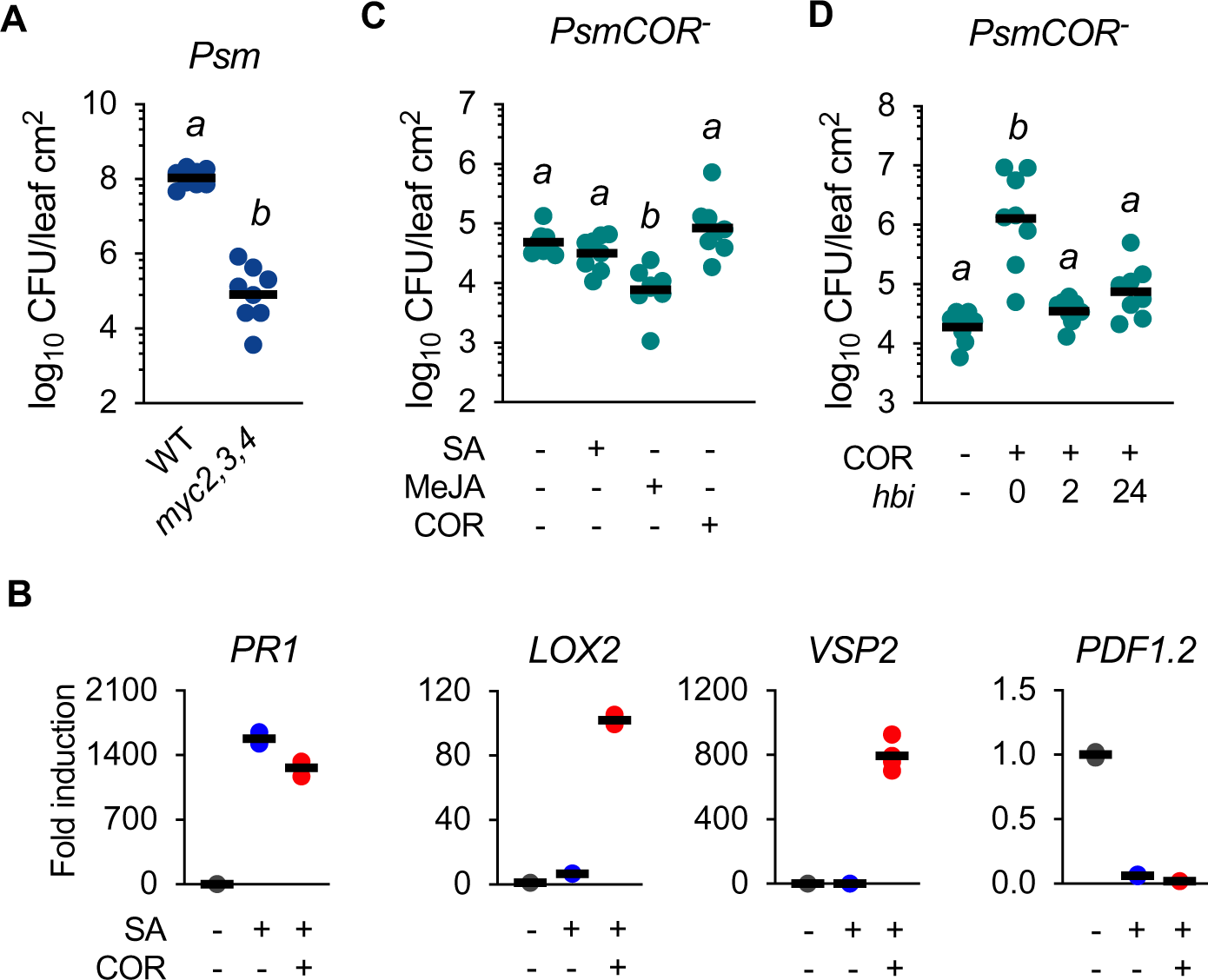
The structural analogues JA and Coronatine induce contrasting immune outcomes. (A) Growth of *Pseudomonas maculicola* ES4326 (*Psm*) in wild-type (WT) and JA-compromised (*myc2,3,4*) plants. Leaves were pressure-inoculated with a bacterial suspension (OD_600_ = 0.005) and growth assessed 3 days after infection. CFU, colony-forming units. n = 8, Tukey ANOVA test, α = 0.05. (B) Transcriptional levels of SA- (*PR1*) and JA- (*LOX2*; *VSP2; PDF1.2*) responsive genes after pharmacological treatment. Transcripts were analysed by semi-quantitative RT-PCR in seedlings treated with 0.5 mM salicylic acid (SA) in the presence or not of 0.1 mM coronatine (COR) mM for 24 hours and normalized to expression of *UBQ5* (n = 4). (C) Growth of coronatine-deficient *Pseudomonas maculicola* ES4326 (*PsmCOR^-^*) in immunized plants. Wild-type plants were immunized with 0.5 mM salicylic acid (SA), 0.1 mM methyl-jasmonate (MeJA) or 0.1 mM coronatine (COR). After 24 hours, leaves were pressure-inoculated with bacterial suspension (OD_600_ = 0.002) and growth assessed 5 days after infection. CFU, colony-forming units. n = 8, Tukey ANOVA test, α = 0.05. (D) Growth of coronatine-deficient *Pseudomonas maculicola* ES4326 (*PsmCOR^-^*) in immunized plants. Wild-type plants were immunized with with 0.1 mM coronatine (COR) 24, 2 or 0 hours before pressure-inoculation (*hbi*) with *PsmCOR^-^* (OD_600_ = 0.002). Bacterial growth was assessed 5 days after infection. CFU, colony-forming units. n = 8, Tukey ANOVA test, α = 0.05.

To further examine the role of COR in *Psm* pathogenicity, we immunized wild-type plants with either SA, JA or COR for 24 hours before inoculation with the non-pathogenic *PsmCOR^-^*. While treatment with SA or COR showed no impact on *PsmCOR^-^* pathogenicity, immunization with JA enhanced plant resistance to this pathogen (Fig. 1C). Because evidence indicate that pathogens hijack the plant hormonal balance early in infection (Katsir et al., 2008; Melotto et al., 2006, 2017), we sought to restore *PsmCOR^-^* pathogenicity by treating plants with COR at earlier time points before infection. While leaf spray with COR for 24 or 2 hours before inoculation showed no impact on *PsmCOR^-^*pathogenicity (Fig. 1D), concomitant bacterial inoculation and leaf spray with COR increased *PsmCOR^-^* growth in wild-type plants by over 30-fold (Fig. 1D). Collectively, these data suggest that, unlike JA, COR functions transiently in immunity by creating an early window of opportunity for pathogenicity.

### Coronatine represses redox-sensitive recruitment of the plant master immune regulator NPR1

The transcriptional coactivator NPR1 is a master regulator of immune responses to *Pseudomonas* (Nomoto et al., 2021). Accordingly, *npr1-1* mutant plants display enhanced disease susceptibility to both *Psm* and *PsmCOR^-^* when compared to wild-type plants (Fig. 2A & B, respectively). To gain further insights on the mechanisms by which COR promotes *Pseudomonas* pathogenicity, we investigated recruitment of NPR1 during infection. Time-course analysis showed that *NPR1* transcription was activated within 4 hours following infection with *PsmCOR^-^*, a response markedly suppressed by application of exogenous COR or upon infection with *Psm* (Fig. 2C). In cells, NPR1 is found in a check oligomer state that, upon infection, is rapidly reduced to its transcriptionally-active, monomeric form to control immune transcriptome (Kneeshaw et al., 2014; Mou et al., 2003). Analysis of NPR1-GFP protein showed that *PsmCOR^-^* elicited accumulation of monomeric NPR1 2 hours after infection, again a response markedly suppressed by exogenous application of COR or upon infection with *Psm* (Fig. 2D). Accordingly, transcription of the NPR1-transcriptinal activity marker gene, *PR1*, was only partially activated upon *Psm* treatment compared to *PsmCOR*^-^ (Fig. 2E). Taken together, these data indicate that COR supresses activation of NPR1 and downstream transcriptional changes early in infection to promote *Pseudomonas* pathogenicity.

**Figure 2.**
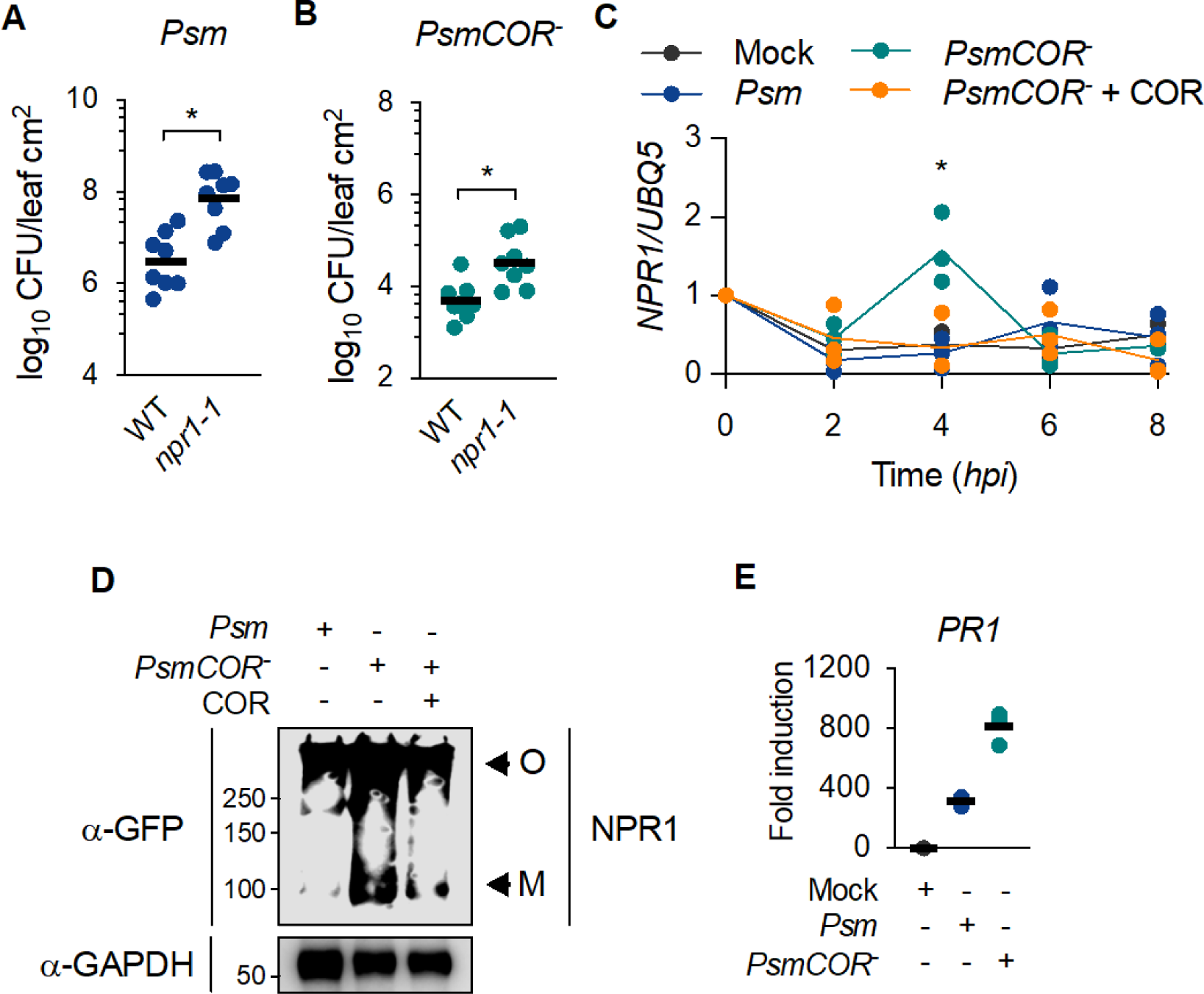
Coronatine represses redox-sensitive recruitment of the plant master immune regulator NPR1. (A) Growth of *Pseudomonas syringae maculicola* ES4326 (*Psm*) in wild-type (WT) and SA-compromised (*npr1-1*) plants. Leaves were pressure-inoculated with a bacterial suspension (OD_600_ = 0.0002) and growth assessed 5 days after infection. CFU, colony-forming units. n = 8, Tukey ANOVA test, α = 0.05. (B) As in (A) but plants were pressure-inoculated with *PsmCOR^-^* (OD_600_ = 0.002). CFU, colony-forming units. n = 8, Tukey ANOVA test, α = 0.05. (C) Transcription levels of *NPR1* in response to *Pseudomonas* infection. Transcripts were analysed by semi-quantitative RT-PCR in wild-type plants leaf sprayed with *Pseudomonas syringae maculicola* ES4326 (*Psm*) or its coronatine-(COR) deficient cognate *PsmCOR^-^* (OD_600_ = 0.002) in the presence or not of exogenous 0.1 mM coronatine (COR) and normalized to expression of *UBQ5* (n = 3, Student’s t-test, α = 0.05). (D) NPR1-GFP conformation in response to *Pseudomonas* infection. Plants expressing NPR1-GFP under the control of the constitutive promoter 35S were leaf sprayed with *Psm* or *PsmCOR^-^* (OD_600_ = 0.002) and oligomer (O) or monomers (M) of NPR1-GFP analysed using anti-GFP antibody 2 hours after treatment, whereas GAPDH shows equal loading. (E) Transcriptional levels of SA-responsive gene *PR1* in response to *Pseudomonas* infection. Transcripts were analysed by semi-quantitative RT-PCR in wild-type plants pressure-infiltrated with *Psm* or *PsmCOR^-^* (OD_600_ = 0.002) for 24 hours and normalized to expression of *UBQ5* (n = 3).

### Coronatine hijacks homeostasis of host redox-sensitive immune regulator GSH

During immune responses, NPR1 conformational changes are driven by fluctuations in the host cellular redox state (Mou et al., 2003; Spoel et al., 2009). Therefore, we examined whether COR impacts levels of the redox-sensitive master regulator of SA immunity, glutathione (GSH), in hosts. Wild-type plants treated with *PsmCOR^-^* displayed higher levels of GSH within 2 hours of infection compared to mock-treated plants, a response abolished or delayed to 4 hours when exogenous COR was applied or plants were infected by *Psm*, respectively (Fig. 3A). Similarly, immunization with SA in the presence or not of JA induced accumulation of total GSH in wild-type plants, a response abolished by COR co-treatment (Fig. 3B). In plants, the F-box protein coronatine-insensitive 1 (COI1) is a master regulator of transcriptional changes in response to JA (Chini et al., 2007). Because COR is thought to function as an agonist of COI1 receptor to promote *Pseudomonas* pathogenicity (Katsir et al., 2008), we assessed GSH accumulation in *coi1-1* knockout mutants. While co-treatment of wild-type plants with SA and COR suppressed GSH levels compared to SA treatment alone (Fig. 3C), *coi1-1* co-treatment with COR showed no impact on GSH levels (Fig. 3C). Collectively, these findings indicate that COR transiently targets host redox signalling to promote *Pseudomonas* pathogenicity.

**Figure 3.**
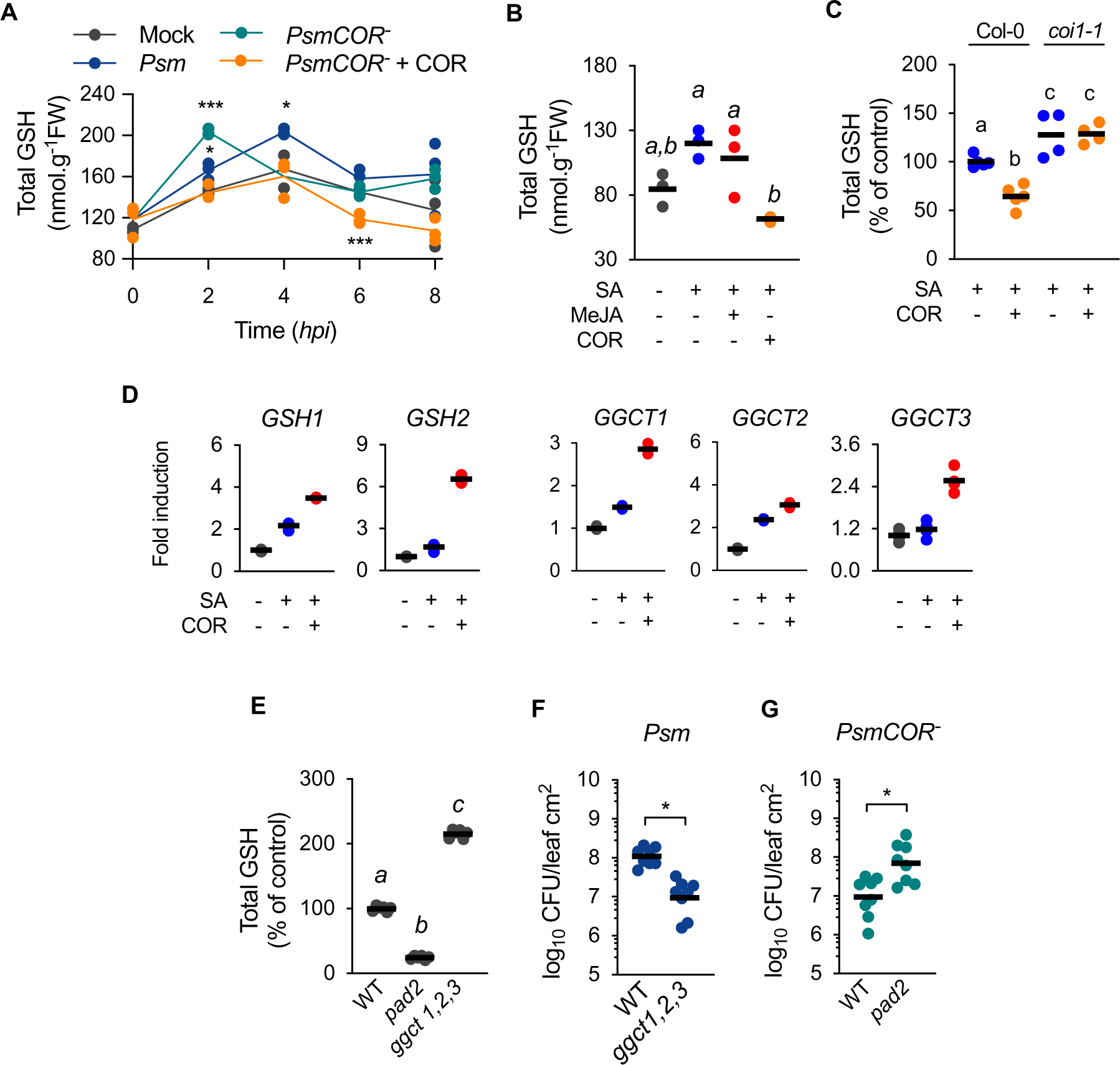
Coronatine hijacks homeostasis of host redox regulator GSH. (A) Levels of the redox-active, immune mediator glutathione (GSH) in response to *Pseudomonas* infection. Wild-type plants were leaf sprayed with *Pseudomonas syringae maculicola* ES4326 (*Psm*) or its coronatine- (COR) deficient cognate *PsmCOR^-^* (OD_600_ = 0.002) in the presence or not of exogenous 0.1 mM COR and leaf total GSH levels assessed at the indicated time (n = 3, Student’s t-test, α = 0.05). (B) As in (A) but wild-type plants were immunized with 0.5 mM salicylic acid (SA) in the presence or not of 0.1 mM methyl-jasmonate (MeJA) or 0.1 mM coronatine (COR). Leaf total GSH levels were estimated after 4 hours of treatment (n = 3, Tukey ANOVA test, α = 0.05). (C) As in (A) but wild-type and JA-compromised *coi1-1* plants were immunized with 0.5 mM salicylic acid (SA) in the presence or not of 0.1 mM coronatine (COR). Leaf total GSH levels were estimated after 4 hours of treatment and shown as percentage of SA for each genotype (n = at least 4, Student’s t-test, α = 0.05). (D) Transcriptional levels of genes committed to glutathione (GSH) synthesis (*GSH1* and *GSH2*) and degradation (*GGCT1*, *GGCT2;1*, *GGCT2;2*) in response to pharmacological treatment. Transcripts were analysed by semi-quantitative RT-PCR in seedlings treated with 0.5 mM salicylic acid (SA) in the presence or not of 0.1 mM coronatine (COR) mM for 24 hours and normalized to expression of *UBQ5* (n = 4). (E) Basal levels of total glutathione (GSH) in GSH mutants *pad2* and *ggct1,2,3*. Leaf total GSH levels were estimated in unchallenged plants (n = 4, Tukey ANOVA test, α = 0.05). (F) Growth of *Psm* in wild-type (WT) and *ggct1,2,3* plants. Leaves were pressure-inoculated with a bacterial suspension (OD_600_ = 0.002) and growth assessed 3 days after infection. CFU, colony-forming units (n = 8, Student’s t-test, α = 0.05). (G) As in (H) but growth of *PsmCOR^-^* was assessed in wild-type (WT) and *pad2* plants. CFU, colony-forming units (n = 8, Student’s t-test, α = 0.05).

To gain insights on the mechanisms by which COR controls GSH homeostasis during immunity, we assessed the impact of SA and COR co-treatments on transcription of genes committed to GSH synthesis (*GSH1* and GSH2) and catabolism (*GGCT1, GGCT2* and *GGCT3*) (Kumar et al., 2015; Ohkama-Ohtsu et al., 2008). When compared to mock-treated plants, transcription of GSH synthetic genes *GSH1* and *GSH2* was activated by SA and COR co-treatment, but not by SA treatment alone. Moreover, while SA treatment of wild-type plants induced minor changes in transcription of GSH catabolic genes compared to mock treatment, co-treatment with COR markedly activated transcription of GSH catabolic genes *GGCT1* and *GGCT3* (Fig. 3D). These data suggest that COR recruits catabolic GGCTs to manipulate GSH homeostasis in hosts. Next, we assessed whether genetic manipulation of basal GSH levels in plants impacts outcome of infection with *Pseudomonas*. Compared to wild-type plants, *pad2* (also known as *gsh1*) and *ggct1,2,3* triple mutants displayed lower and higher basal levels of GSH, respectively (Fig. 3E). Accordingly, while *ggct1,2,3* mutants displayed enhanced disease resistance to *Psm* (Fig. 3F), *pad2* plants displayed enhanced susceptibility to *PsmCOR^-^* (Fig. 3G), suggesting that reduced levels of GSH in hosts complement *PsmCOR^-^* weak pathogenicity phenotype. Overall, these data show that COR-induced pathogenicity is underpinned by changes in host GSH homeostasis.

### Coronatine promotes *Pseudomonas* pathogenicity by driving specificity in protein-SNO signalling

In cells, GSH can spontaneously react with nitric oxide (NO) to form the *trans*-nitrosylating agent, S-nitrosoglutathione (GSNO) (Feechan et al., 2005; Liu et al., 2001). Although both, free NO and GSNO, can modify proteins post-translationally to form *S*-nitrosothiols (protein-SNO) (Frungillo et al., 2014; Kneeshaw et al., 2014), these *S-*nitrosylating agents define genetically distinct branches of NO signalling in plants (Kneeshaw et al., 2014; Yun et al., 2016). To assess whether COR drives specificity in protein-SNO signalling to promote *Pseudomonas* pathogenicity, we took advantage of mutant plants that accumulate excessive amounts of protein-SNO due to over accumulation of GSNO or over production of NO, *gsnor1* (also known as *par2-1*) and *nox1* (Chen et al., 2009; Feechan et al., 2005; He et al., 2004), respectively. As reported previously (Kneeshaw et al., 2014; Yun et al., 2011), both *nox1* and *gsnor1* mutant plants displayed enhanced disease susceptibility to pathogenic *Psm*, showing over 20-fold higher bacterial growth compared to wild-type plants (Fig. 4A). Interestingly, whereas *gsnor1* mutants displayed resistance comparable to wild-type, *nox1* showed marked susceptibility to *PsmCOR^-^* (Fig. 4B), indicating that over production of NO in hosts is sufficient to rescue *PsmCOR^-^* pathogenicity.

**Figure 4.**
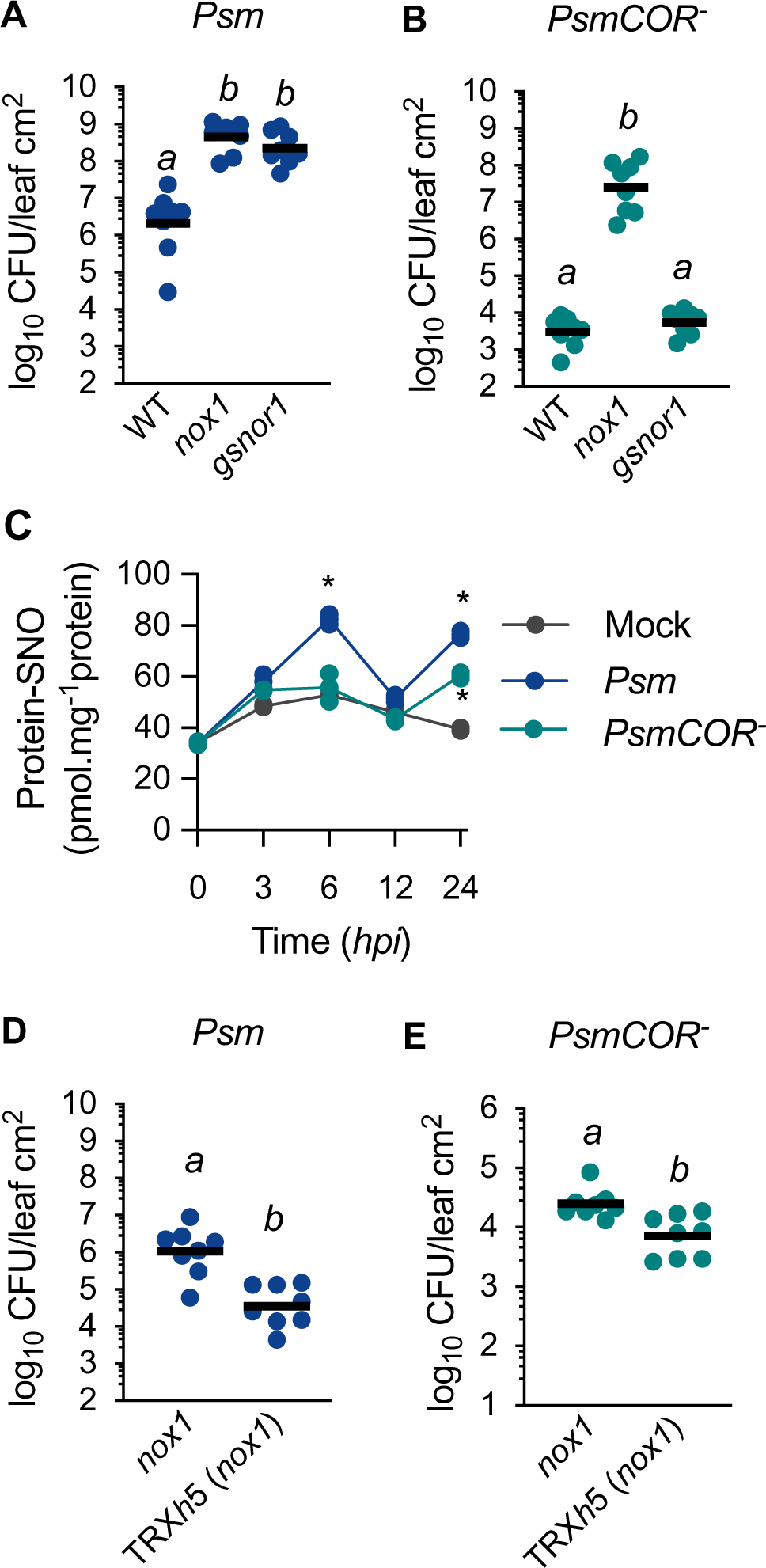
Coronatine promotes *Pseudomonas* pathogenicity by driving specificity in protein-SNO signalling. (A) Growth of *Pseudomonas maculicola* ES4326 (*Psm*) in wild-type (WT) and SNO mutants (*nox1* and *gsnor1*). Leaves were pressure-inoculated with a bacterial suspension (OD_600_ = 0.0002) and growth assessed 5 days after infection. CFU, colony-forming units. n = 8, Tukey ANOVA test, α = 0.05. (B) As in (A) but growth of *PsmCOR^-^*(OD_600_ = 0.002) was assessed. CFU, colony-forming units. n = 8, Tukey ANOVA test, α = 0.05. (C) Total *S-*nitrosylated protein content (protein-SNO) was estimated in leaf extracts of wild-type plants sprayed with virulent *Psm* or avirulent *PsmCOR^-^* in the indicated time points (n = 3). Asterisk indicate statistically significant differences for each treatment point compared to its respective control (n = 3, one-way ANOVA test, *P < 0.05, *P < 0.01). (D) As in (A) but growth of *Psm* (OD_600_ = 0.0002) was assessed in *nox1* and cognate TRX*h*5 overexpressing line. CFU, colony-forming units. n = 8, Tukey ANOVA test, α = 0.05. (E) As in (A) but growth of *PsmCOR^-^*(OD_600_ = 0.002) was assessed in *nox1* and cognate TRX*h*5 overexpressing line. CFU, colony-forming units. n = 8, Tukey ANOVA test, α = 0.05.

To further investigate if specificity in protein-SNO confers pathogenicity to *Pseudomonas*, we assessed protein-SNO levels over time in wild-type plants challenged with *Psm* or *PsmCOR^-^*. As expected, compared to control, bacterial infection induced accumulation of protein-SNO in both *Psm*- or *PsmCOR^-^*-treated plants after 24 hours (Fig. 4C). Remarkably, infection by *Psm*, but not *PsmCOR^-^*, resulted in higher levels of protein-SNO after 6 hours of treatment (Fig. 4C), corroborating the notion that COR targets protein-NO signalling in plants. Control of protein-SNO levels and downstream signalling in *nox1* is achieved through the selective activity of the protein-SNO reductase, Thioredoxin *h*5 (TRX*h*5) (Kneeshaw et al., 2014). We, therefore, investigated if constitutive expression of TRX*h*5 rescues *nox1* susceptibility phenotype. Strikingly, overexpression of TRX*h*5, restored *nox1* resistance to pathogenic *Psm* and non-pathogenic *Pseudomonas* strains (Fig. 4D and E, respectively). Taken together, these data indicate COR induces a drift between specific branches of protein-SNO signalling in plants to promote *Pseudomonas* pathogenicity.

## DISCUSSION

Reprogramming of the redox metabolism underlies plant immune responses to microbial pathogens. To fine tune the immune response according to attackers’ invading strategy, plants have evolved sophisticated signalling networks based on complex relationships between phytohormones (Pieterse et al., 2009; Spoel and Dong, 2012; Tsuda and Somssich, 2015). Compensatory interactions between major signalling sectors in the plant immune network are thought to confer immune resilience and attenuate interferences mediated by attackers (Hillmer et al., 2017; Spoel et al., 2009; Tsuda et al., 2009). Pathogens, however, often succeed in infection, suggesting that hubs in the plant immune network are hijacked. Our study on the phytotoxin COR shows that adapted pathogens promote virulence by offsetting homeostasis of reactive nitrogen intermediates in host to effectively evade immunity.

We show that, while sharing signalling components in plants, the phytohormone JA and the phytotoxin COR induce contrasting immune outcomes. Plant immune responses against the bacterial pathogen *Pseudomonas* is largely mediated by the phytohormone SA (Pieterse et al., 2009; Spoel and Dong, 2012). Although at high concentrations antagonistic effects have been extensively reported, at low concentrations the immune phytohormones SA and JA have transient synergistic effects on transcription of immune genes (Mur et al., 2006). Our findings add to the complexity of plant immune network by demonstrating that immunization with JA, but not SA, induces further resistance against *PsmCOR^-^* (Fig. 1). JA binding to the ubiquitin-ligase SCF^COI1^ complex triggers degradation of JAZ transcriptional suppressors by the proteasome pathway, relieving the inhibition on MYC transcription factors and downstream gene expression (Chini et al., 2007; Fernández-Calvo et al., 2011; Laurie-Berry et al., 2006). The MYC family of transcription factors mediates expression of a set of JA-responsive genes (e.g. *VSP2*), while antagonizing alternative transcriptional responses mediated by JA (e.g. *PDF1.2*) (Fernández-Calvo et al., 2011; Lorenzo et al., 2004). Because JA and COR treatments induced distinct transcriptional responses during immunity (Fig. 1), we propose that trade-offs within JA signalling allow plants to exploit *Psm* attack strategy to orchestrate immune responses. The fact that the chemical structure of COI1 ligand underpins its selectivity towards JAZ transcriptional repressors (Campos et al., 2016; Takaoka et al., 2018; Zhang et al., 2015) could provide an explanation to the distinct immune outcomes observed in this study.

Our work shows that specificity in redox signalling underlies immune outcome in plants, a trait exploited by pathogens to promote disease. We discovered that COR transiently prevents early GSH accumulation upon infection to promote bacterial pathogenicity (Fig. 2). These findings coincide with the time window for immune hormonal cross talk (Koornneef et al., 2008), indicating that timely manipulation of the host intracellular redox environment confers *Pseudomonas* with advantage over the plant immune network. The findings that immunization with SA triggered fast accumulation of GSH, despite minor effects on transcription of genes committed to GSH synthesis (Fig. 3) suggest that a metabolic shift controls GSH levels during early stages of infection. Synthesis of GSH is achieved by the conjugation of GLU, CYS and GLY in a two ATP-dependent steps (Noctor and Foyer, 1998). Condensation of GLY with γ-glutamyl-cysteine (γ-EC), the terminal step in GSH synthesis, plays a critical role in GSH synthesis and accumulation (Noctor et al., 1997, 1999). Under CO_2_ deficit, GLY accumulates through photorespiration, a process aimed at minimizing the deleterious effect of surplus in reducing power generated by electron flow in the photosystem (Noctor et al., 1997, 1999). Intriguingly, COR quickly reverts stomatal closure induced by microbial associated molecular pattern (MAMP), an immune response thought to reduce infection rate by physically restricting microbial access to plant tissue (Melotto et al., 2006, 2017). Because stomatal closure restricts gas exchange, it is plausible that rapid GSH accumulation is favoured by photorespiratory GLY induced at low CO_2_ availability upon MAMP recognition by hosts. In agreement, low CO_2_ atmosphere has been shown to enhance disease resistance of *Arabidopsis* to *Pseudomonas* independently of stomatal movement (Zhou et al., 2017). Thus, we propose that, besides controlling infection rate, stomatal movement signals for a metabolic shift that shapes intracellular redox environment to tune immune response.

Conformation and stability of the master transcriptional regulator NPR1 is controlled by the redox-based post-translational modification S-nitrosylation (Kneeshaw et al., 2014; Mou et al., 2003; Spoel et al., 2009; Tada et al., 2008). Upon bacterial infection, we found that COR supresses accumulation of active NPR1 (Fig. 2). Additionally, unbalanced NO signalling and protein-SNO levels in *nox1* and *gsnor1* mutant plants are associated with enhanced susceptibility to *Psm* (Fig. 4) (Feechan et al., 2005; Yun et al., 2011, 2016). We found that, in contrast to *gsnor1*, over production of NO in *nox1* mutants restored pathogenicity of *PsmCOR^-^*, a phenotype reversed by expression of the protein-SNO reductase TRX*h*5 (Fig. 4). These data demonstrate that *Pseudomonas* use COR to drive specificity in protein-SNO signalling in hosts as a mechanism of pathogenicity.

In conclusion, our findings demonstrate that *Pseudomonas* disarms the plant immune network by driving specificity in redox signalling in hosts within an early time window following infection. In addition to its established role as a major antioxidant in cells, we demonstrate that GSH pool determines specificity in NO-based post-translational modifications during immunity and propose that protein-SNO signalling underlies trade-offs in plant hormonal immune networks. Targeting the chemical equilibrium between reactive nitrogen intermediates may prove a fruitful strategy for engineering complex signalling networks in higher plants.

## ACKNOWLEDGEMENTS

This work was supported by the European Molecular Biology Organization (EMBO, DE) grant 420-2015 (to L.F.), by Biotechnology and Biological Sciences Research Council (BBSRC) grant BB/S010262/1 (to L.F.), by European Research Council (ERC) under the European Union’s Horizon 2020 research and innovation program, grant agreements no. 678511 and no. 101001137 (to S.H.S.), by Royal Society University Research Fellowships no. UF090321 and UF140600 (to S.H.S.) and a Royal Society International Exchanges grant IEC\R3\170118 (to S.H.S. and Y.T.).

## AUTHOR CONTRIBUTIONS

LF and SHS designed the research. LF performed experiments. NO, MN, and YT provided crucial research materials. SL and BY performed chemiluminescence assays. LF, YT, BY and SHS performed project administration and acquired funding. The manuscript was written by LF and SHS.

## MATERIAL AND METHODS

### Plant material and growth conditions

*Arabidopsis thaliana* ecotype Columbia-0 (WT), cognates mutant lines *npr1-0* (Alonso et al., 2003)*; npr1-1* (Cao et al., 1994)*; pad2* (Parisy et al., 2007)*; coi1-1* (Xie et al., 1998)*; myc2,3,4* (Fernández-Calvo et al., 2011)*; ggct1,2,3; nox1* (He et al., 2004); *gsnor1* (*par2-1*) (Chen et al., 2009) and transgenic line 35S::TRX*h*5 (Kneeshaw et al., 2014) were grown in soil at 21°C and 100 μmoL·m^−2^·s^−1^ light, 50-65% relative humidity and a photoperiod of 16/8 h light/dark. For experiments on seedlings, seeds were surface sterilized with 50% bleach, 1% Tween-20 for 15 min, washed five times with sterile water and sown aseptically in petri dishes containing MS medium. Seedlings were cultivated as described above. Twelve-day-old seedlings were used for the experiments.

### Oxidative burst assay

Generation of reactive oxygen species (ROS) in response to synthetic MAMP flagellin peptide flg22 was estimated by chemiluminescence as described elsewhere (Smith and Heese, 2014). Leaf discs were placed in 96-well plates and incubated overnight as indicated in the figure legends. After removal of the aqueous solution, ROS production was elicited by incubation with 0.1 µM flg22 in the presence of 200 ng/mL luminol and 20 µg/mL horseradish peroxidase. Chemiluminescence measurements were taken using a plate reader (Infinite 200 PRO, Tecan, Switzerland) every 2.5 min for 2 hours.

### Pathogen infection and chemical induction

For pathogen infection, *Pseudomonas syringae* pv. *maculicola* (*Psm*) ES4326 and cognate, coronatine-deficient *PsmCOR^-^* were grown overnight in liquid LB medium supplemented with 10 mM MgSO_4_ and streptomycin (100 μg/mL) or kanamycin (50 μg/mL), respectively. Bacterial suspensions in 10 mM MgSO_4_ were pressure infiltrated via the abaxial side of leaves at concentration indicated in the figure legends. *In planta* bacterial growth was determined by spreading serial dilutions of leaf extracts on LB plates supplemented with 10 mM MgSO_4_, 100 μM cycloheximide and antibiotics as appropriated.

For chemical induction of immune responses, plants were leaf sprayed with aqueous solution containing 0.5 mM sodium salicylate (SA), 0.1 mM Methyl-Jasmonate (MeJA) or 0.1 mM coronaine (COR) 24 hours before infection, unless otherwise stated in the figure legends. For the analysis of biochemical parameters, plants were leaf sprayed with bacterial suspension at dawn. To this end, bacterial cells were grown as described above and diluted in 10 mM MgSO_4_, 0.02% Silwet L-77.

### Gene expression analysis

For real-time PCR analysis, RNA extraction and cDNA synthesis were performed as described previously (Spoel et al., 2009). Total RNA was extracted in phenol:chlorophorm mixture and used as template for cDNA synthesis using ImProm II reverse transcriptase, as recommended by the manufacturer. Gene expression analysis was carried out using SYBR Green Master Mix in a Real-Time PCR System 7500 (Applied Biosystem, Foster City, CA) and calculated using the 2^-ΔDDCt^ method (Livak and Schmittgen, 2001) with *UBQ5* as internal standard (Kneeshaw et al., 2014).

### Protein analysis

Leaves were mechanically grounded and resuspended in 50 mM Tris-HCl, pH 7.5, 5mM EDTA; 150 mM NaCl, 10% glycerol and proteinase inhibitors (50 mg ml^-1^ TPCK; 50 mg ml^-1^ TLCK; 0.5 mM PMSF). After centrifugation at 14,000*xg* at 4°C, protein extracts were separated on an 8% SDS-PAGE gel and transferred onto a nitrocellulose membrane. Blots were probed with anti-GFP (1:2,500) or anti-GAPDH (1:2,000) and anti-rabbit or anti-mouse conjugated with horseradish peroxidase antibody (1:1,000) diluted in blocking buffer (1x PBS containing 5% non-fat milk and 0.1% Tween 20). Antibody-bound proteins were detected by using SuperSignal West Pico PLUS chemiluminescent substrate.

### Total glutathione estimation

Total leaf glutathione (GSH + GSSG) content was estimated spectrophotometrically as the rate TNB production as described elsewhere (Rahman et al., 2007). Leaves were harvested following indicated treatment and frozen immediately in liquid N_2_.

Tissue samples were mechanically homogenized and resuspended (1:10 w/v) in 0.1 M potassium phosphate buffer, pH 7.5, containing 1 mM EDTA, 30 mM 5-sulfosalicylic acid, 0.1% Triton X-10. After centrifugation at 3,000*xg* for 5 min at 4°C, aliquots of the supernatant were incubated in 0.1 M potassium phosphate buffer, pH 7.5, containing 1 mM EDTA, 0.6 mM 5,5′-dithio-bis(2-nitrobenzoic acid), 0.1 mM NADPH, 5 U/mL of yeast glutathione reductase. Absorbance measurements were taken at 412 mm (Infinite 200 PRO, Tecan, Switzerland) every 35 sec for 3 min. The obtained values were compared against those of a standard curve constructed using GSH and normalized by fresh weight. All samples were tested for linearity.

### Protein-SNO determination

Protein-SNO levels in leaf extracts were determined by chemiluminescence as described previously (Yun et al., 2011). Protein homogenates were purified in G-25 fine Sephadex chromatography column equilibrated with 1x PBS, pH7.4, 0.5 mM EDTA. After reaction with ozone, NO liberated from nitrosyl groups by HgCl_2_ were measured using a Sievers nitric oxide analyser. The obtained values were compared against a standard curve constructed using GSNO and normalized by protein content determined by Coomassie-blue method, as recommended by the manufacturer. All samples were protected from light during the assay.

### Statistical analysis

Statistical analysis used and sample size (n) are indicated in the figure legends. For all data, a minimum of two experiments with similar results were performed.

